# Multitask deep learning for the emulation and calibration of an agent-based malaria transmission model

**DOI:** 10.1101/2025.02.05.636692

**Authors:** Agastya Mondal, Rushil Anirudh, Prashanth Selvaraj

## Abstract

Agent-based models of malaria transmission are useful tools for understanding disease dynamics and planning interventions, but they can be computationally intensive to calibrate. We present a multitask deep learning approach for emulating and calibrating a complex agent-based model of malaria transmission. Our neural network emulator was trained on a large suite of simulations from the EMOD malaria model, an agent-based model of malaria transmission dynamics, capturing relationships between immunological parameters and epidemiological outcomes such as age-stratified incidence and prevalence across eight sub-Saharan African study sites. We then use the trained emulator in conjunction with parameter estimation techniques to calibrate the underlying model to reference data. Taken together, this analysis shows the potential of machine learning-guided emulator design for complex scientific processes and their comparison to field data.

**Author summary:** Mathematical models can help understand and design interventions for infectious diseases such as malaria, but they can be complex and time-consuming to work with. In this study, we developed a machine learning approach to create a fast and accurate emulator for a detailed malaria transmission model. Our emulator can quickly predict various disease outcomes based on different immune system parameters. We trained it using a large set of simulations created using the EMOD malaria model and tested it against real-world data from multiple African countries. The emulator not only matches the original model’s predictions closely, but can also be used to efficiently find the best parameter values to match field observations. We hope that this work can serve as a demonstration of machine learning tools to link complex models of disease transmission to field data.

## Introduction

Despite progress in recent decades, the burden of malaria remains unacceptably high, especially in sub-Saharan Africa [1]. Due to numerous factors including seasonal heterogeneity, complex immune landscapes, and the spatial distribution of vectors and interventions, designing effective malaria control strategies is difficult. Mathematical models of disease transmission can aid in the design of these programs by parameterizing the scientific processes that underlie transmission in a given setting. For novel interventions such as genetic vector control tools and updated vaccines [2, 3], for which field data do not yet exist, mathematical models can bridge the gap between expected outcomes and intervention parameters. Epidemiological models have been widely used for policy planning, prediction, and etiology [4–8]. These models generally provide a mechanistic framework by which to understand the progression of a disease from biological to sociodemographic facets.

For vector-borne diseases such as malaria, the progression of disease represents a complex relationship between vector, host, and environment. Although simple models of malaria transmission have been used for more than a century [9], in recent years more complex models have been developed. These models explicitly parameterize the development of the pathogen in the vector and can take into account heterogeneity in transmission such as age structure, immunity, species, and spatial landscape. While these models can accurately describe the processes involved in disease progression, they tend to require large amounts of computational resources to estimate parameters for a wide range of outcomes and settings. From a biological perspective, the reciprocal transmission of the malaria pathogen between vectors and humans is a detailed process involving vector feeding behavior, the development of infectious gametocytes in human liver and blood stages, and various forms of anti-parasite immunity. Modeling malaria transmission in large populations additionally requires attention to environmental conditions (as rainfall is a major ecological driver of larval carrying capacity), vector control interventions, mobility, and vector species. Clinical manifestations of malaria add another layer of complexity, whereby humans can be infected clinically, asymptomatically, or subpatently. Thus, models of malaria transmission tend to be more complex than those of other disease systems, such as respiratory viruses. Simple models of malaria transmission include numerous simplifying assumptions. For example, the Ross-Macdonald model [9] (perhaps the simplest malaria transmission model), represents humans and vectors as homogeneous, randomly-mixing populations, and considers only susceptible and infectious states. More complex models such as EMOD [10, 11] and OpenMalaria [12] explicitly describe the emergence of infection in stochastic, individual-based, formulations. Evaluating complex models for large sets of scenarios therefore requires sufficient computational resources.

Additionally, calibrating (i.e., aligning model parameters) to external field data can be challenging, as exploring multidimensional parameter spaces can quickly become intractable. These techniques therefore demand large amounts of computational resources. For complex systems such as mechanistic disease transmission models, parameter estimation techniques such as Bayesian Markov chain Monte Carlo, sequential Monte Carlo, and approximate Bayesian computation have been employed to calibrate mechanistic models [13–16]. Resource-constrained methods such as Bayesian likelihood calculations [17] and nearest neighbors [18] allow for parameter estimation with respect to complex models, but can be limited by sampling and defining explicit likelihood structures.

Recent literature has proposed the development of machine learning (ML) model “surrogates” or “emulators” to complex disease transmission models [18–20]. These models allow for the computationally efficient evaluation and prediction of disease dynamics as compared to the original model, and generally learn patterns from a large set of realizations from the underlying model. While there are many techniques to develop an emulator model [21–23], we focus here on the application of deep learning as an emulator for a complex model of malaria transmission. Deep neural networks (DNNs) represent a class of models that enable the estimation of an arbitrary function between inputs and outputs. Unlike other emulators, DNNs make very few assumptions about the functional relationship between the input and output space, and are flexible and extensible to a large variety of use cases. Additionally, once trained, evaluation of a DNN is orders of magnitude faster than a realization of the underlying mechanistic model. Finally, DNNs were chosen for this analysis because, unlike other methods, DNNs are known to model stochastic processes that are important to consider in disease transmission, as infection events can be governed by an underlying probability distribution. Incorporating stochastic realizations into the training of the DNN emulator allows for the inclusion of stochastic uncertainty, as the emulator captures both the mean behavior and the spread across runs.

Mathematically, we are interested in training a deep neural network to infer a function *f* mapping a vector of parameters **x** to a set of prevalence and incidence outputs *y_i_*, for *i* ∈ [1*, n*] outputs, where the inputs and outputs correspond to realizations of the original mechanistic model:

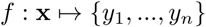

Each input parameter set will map to multiple prevalence and incidence outputs simultaneously, leveraging methods from multitask learning. In this framework, one neural network predicts multiple outcomes, allowing the network to learn shared patterns in the simulation data. The need to train several emulators separately is additionally eliminated [24].

With respect to calibration, the trained emulator can be used to rapidly search parameter space to identify input immune parameters that result in prevalence and incidence outputs that align with reference data, as the evaluation of trained DNNs orders of magnitude faster than the underlying mechanistic model. Techniques from numerical optimization such as gradient descent allow for more flexibility in exploring parameter space and can identify out-of-distribution calibration fits. We couple these parameter estimation techniques with the trained emulator to identify parameter sets whose outputs most closely align with reference data. Finally, as we are interested in the universality of these biologically relevant model parameters across study sites, supplementary analyses were conducted to test the emulator-calibrated parameter sets against outputs from study sites not seen during training. Thus, in this work, we show that i) deep learning techniques can act as emulators to complex epidemiological models, ii) these deep learning emulators can be used to calibrate the underlying mechanistic model to field data, and iii) model parameters identified by the emulator can capture immune dynamics in study sites not previously seen during training, allowing for inference on new sites without retraining the emulator.

## Methods

### Malaria transmission model and parameter space

Simulations were conducted using the EMOD framework [10, 11]. EMOD is an agent-based, stochastic mechanistic model of malaria transmission that incorporates within-host parasite and immune dynamics alongside vector life cycle dynamics and human demographics. Here, we are interested in calibrating parameters associated with adaptive immunity that is stimulated by blood-stage malaria infection within the human host. Our ultimate goal in this study is to identify a universal set of immune factors whose simulated outputs closely align with collected parasite prevalence and clinical incidence data across eight sub-Saharan African study sites. As such, in order to build a suite of simulations on which the DNN was trained, we generated 2000 Latin hypercube-sampled parameter sets per site, with 10 stochastic repetitions per set, resulting in a total of 160,000 (8 sites x 2000 parameter sets x 10 stochastic repetitions) simulations. Descriptions of the immune parameters and their biologically relevant ranges are given in Table 1. Following the methodology in Selvaraj *et al.* [25], each study site has its own set of entomological innoculation rates (EIRs), antimalarial interventions, and case management parameters, which remain unchanged across simulations for a given site. As outputs, we tracked four types of data corresponding to available reference data: annual clinical incidence by age group, annual *P. falciparum* prevalence by age group, asexual parasite density by age group by month, and gametocyte density by age group by month. Each site’s reference data corresponds to a unique set of age and density bins, and the output for each simulation for a specific site is binned in alignment with the reference data.

**Table 1.** Overview of immune parameters used to generate Latin hypercube samples with which to simulate.

| Parameter | Description | Range |
| --- | --- | --- |
| Antigen switch rate | The antigenic switching rate per infected red blood cell per asexual cycle. | $[10^{-10}, 10^{-9}]$ |
| Falciparum MSP variants | The number of distinct merozoite surface protein variants for *P. falciparum* malaria in the overall parasite population in the simulation, not necessarily in an individual. | $[5, 50]$ |
| Falciparum nonspecific types | The number of distinct non-specific types of *P. falciparum* malaria. | $[5, 100]$ |
| Falciparum PfEMP1 variants | The number of distinct *Plasmodium falciparum* erythrocyte membrane protein 1 (PfEMP1) variants for *P. falciparum* malaria in the overall parasite population in the simulation. | $[600, 1700]$ |
| Max individual infections | The limit on the number of infections that an individual can have simultaneously. | $[3, 10]$ |
| MSP merozoite kill fraction | The fraction of merozoites inhibited from invading new erythrocytes when MSP1-specific antibody level is 1. | $[0.20, 0.80]$ |
| Nonspecific antigenicity factor | The nonspecific antigenicity factor that adjusts antibody iRBC kill rate to account for iRBCs caused by antibody responses to antigenically weak surface proteins. | $[0.1, 0.9]$ |

### Data sources

In order to validate the ability of the model emulator to produce epidemiologically relevant outputs, we calibrated the underlying model against field data from eight sub-Saharan African sites, using the trained emulator. The Garki project [26] was a multi-year study conducted in Nigeria in the 1970s with the goal of understanding the feasibility of malaria elimination in this setting, aided by the collection of parasitology data to supplement epidemiological and entomological measurements. Additional field data from Senegal, Tanzania, and Burkina Faso were used to evaluate the emulator [27–29]. Each reference site recorded data in a structure unique to the site’s resources. The Garki sites (Matsari, Rafin Marke, and Sugungum) recorded month and age-binned asexual parasite density, the Senegal (Ndiop and Dielmo) sites recorded age-binned annual clinical malaria incidence, the Burkina Faso sites (Dapelogo and Laye) recorded month and age-binned asexual parasite and gametocyte density, and the Tanazania site (Namawala) recorded age-binned malaria prevalence. With respect to data collection, parasite density data from the Garki and Burkina Faso sites were collected via microscopy and real-time quantitative nucleic acid sequence-based amplification (QT-NASBA), respectively [26]. Giemsa stains were used to ascertain malaria prevalence in the Tanzania site [27], and active surveillance was used to ascertain clinical malaria incidence in the Senegal sites [28]. Our simulations however record all of these outcomes for each site-specific simulation. In a simulation, we are able to export fine-grained data for a range of outcomes and aggregate them to match the study site reference data. Then, despite a small reference dataset, we can train our emulator on multiple outcomes such that it can infer patterns between the input parameters and each outcome. In the calibration step, when comparing the emulator’s outputs to the reference data for a site, only the relevant outputs are selected and compared. Table 2 shows an overview of the associated data for each study site.

**Table 2.**
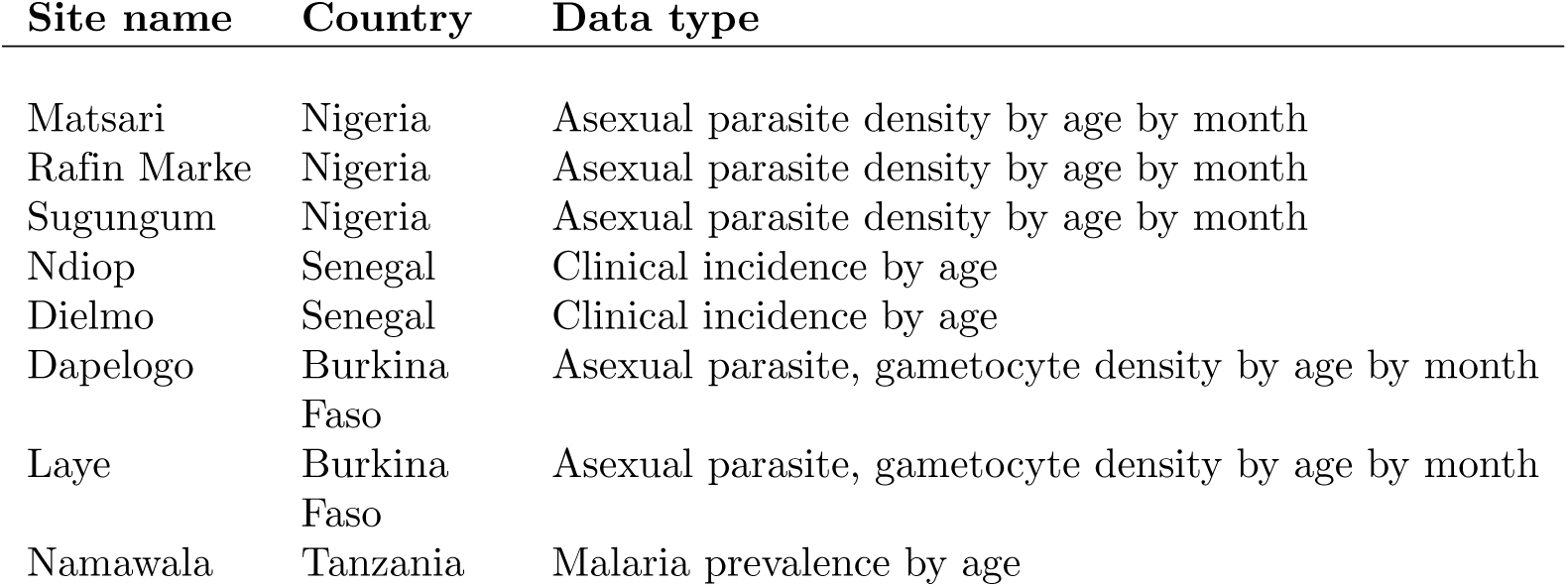
Overview of study sites and reference data to which the model emulator is calibrated.

| Site name | Country | Data type |
| --- | --- | --- |
| Matsari | Nigeria | Asexual parasite density by age by month |
| Rafin Marke | Nigeria | Asexual parasite density by age by month |
| Sugungum | Nigeria | Asexual parasite density by age by month |
| Ndiop | Senegal | Clinical incidence by age |
| Dielmo | Senegal | Clinical incidence by age |
| Dapelogo | Burkina Faso | Asexual parasite, gametocyte density by age by month |
| Laye | Burkina Faso | Asexual parasite, gametocyte density by age by month |
| Namawala | Tanzania | Malaria prevalence by age |

### Deep learning architecture

Once the simulation suite had been run, a DNN was trained on top of simulation outputs. As stated, our goal is to learn a function *f* that maps the input space **x** (the immune parameters and the study site) to the incidence and prevalence outputs *y* for each of the *n* outcomes:

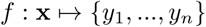

We model *f* as a feed-forward, multitask neural network with six fully-connected shared, hidden layers and five fully-connected task-specific layers. Rectified linear units (ReLU) were used to introduce nonlinearity into the network, and dropout and batch normalization were used to prevent overfitting. Input features were standardized using min-max scaling and outputs were normalized between 0-1 via a sigmoid transformation. The neural network emulator was trained by inputting ten stochastic repetitions of EMOD model outputs for each parameter set, essentially capturing the mean response across simulation runs. While we did not model simulation uncertainty explicitly, neural network architectures such as Bayesian deep convolutional networks [18, 30, 31], allow for the quantification of uncertainty. Explicitly modeling uncertainty may be beneficial in other scenarios, such as sensitivity analyses and the development of model-informed target product profiles [2, 32].

### Model training and hyperparameter tuning

Training a neural network involves iteratively optimizing the model’s “weights” (i.e., the strengths of connection between different layers in the neural network) in order to minimize the discrepancy between model output and training data. Then, the model is evaluated on a previously-unseen data subset to evaluate its performance on new values.

We define the *loss function* as the numerical discrepancy between the neural network’s output and the training data. While there are many loss functions for different modeling goals, a widely-used loss function is the *ℓ*_2_ loss, corresponding to the mean squared error between model outputs *f* (**x**) and training data *y*(**x**) for a given set of input parameters **x** and *N* data samples:

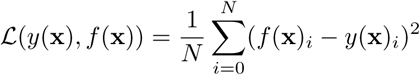

In a multitask learning framework, we further segment this loss function per task, such that the overall goal of training is to minimize the joint loss across *n* tasks and weight *λ_i_*for task *i*:

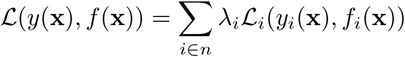

Because our emulator is relatively simple, we employ uniform loss weighting (i.e., *λ_i_* = 1∀*i*), but for more complex networks, the relative importance of each task’s loss becomes important [33]. Many methods have been proposed to address balancing the losses between tasks to ensure the model jointly learns each output type. We use the Adam stochastic gradient descent algorithm [34] to optimize the emulator’s weights and minimize the multitask loss function.

A final component of model specification involves the selection of hyperparameters. In contrast to model weights, which are learned during the optimization process, hyperparameters are fixed attributes of the model that are selected prior to training. These include parameters associated with network architecture (number of hidden layers, number of neurons per layer), training (learning rate, batch size), and regularization (dropout probability, weight decay). In order to select hyperparameters for our model, we randomly sample 100 hyperparameter sets and use the ASHA [35] algorithm to efficiently select the sets that minimize the joint loss on a hold-out data subset after training. Once the neural network is trained, we are able to evaluate its ability to calibrate the underlying model to reference data.

### Calibration workflow

To evaluate the emulator’s performance in identifying the optimal input parameter set for the underlying mechanistic model, outputs from the trained emulator were compared to the reference data to determine the input parameter set whose associated emulated outputs most closely aligned with the reference data. Drawing from techniques in numerical optimization, stochastic gradient descent (SGD) was chosen as the primary parameter estimation method. In this framework, parameter sets were chosen by optimizing against a loss function comparing emulated outputs to the reference data. Iterative updates were made to the input parameters to minimize the loss function until a stopping criterion was achieved. Fig 1 shows a schematic workflow of the emulation and parameter estimation process.

**Fig 1.**
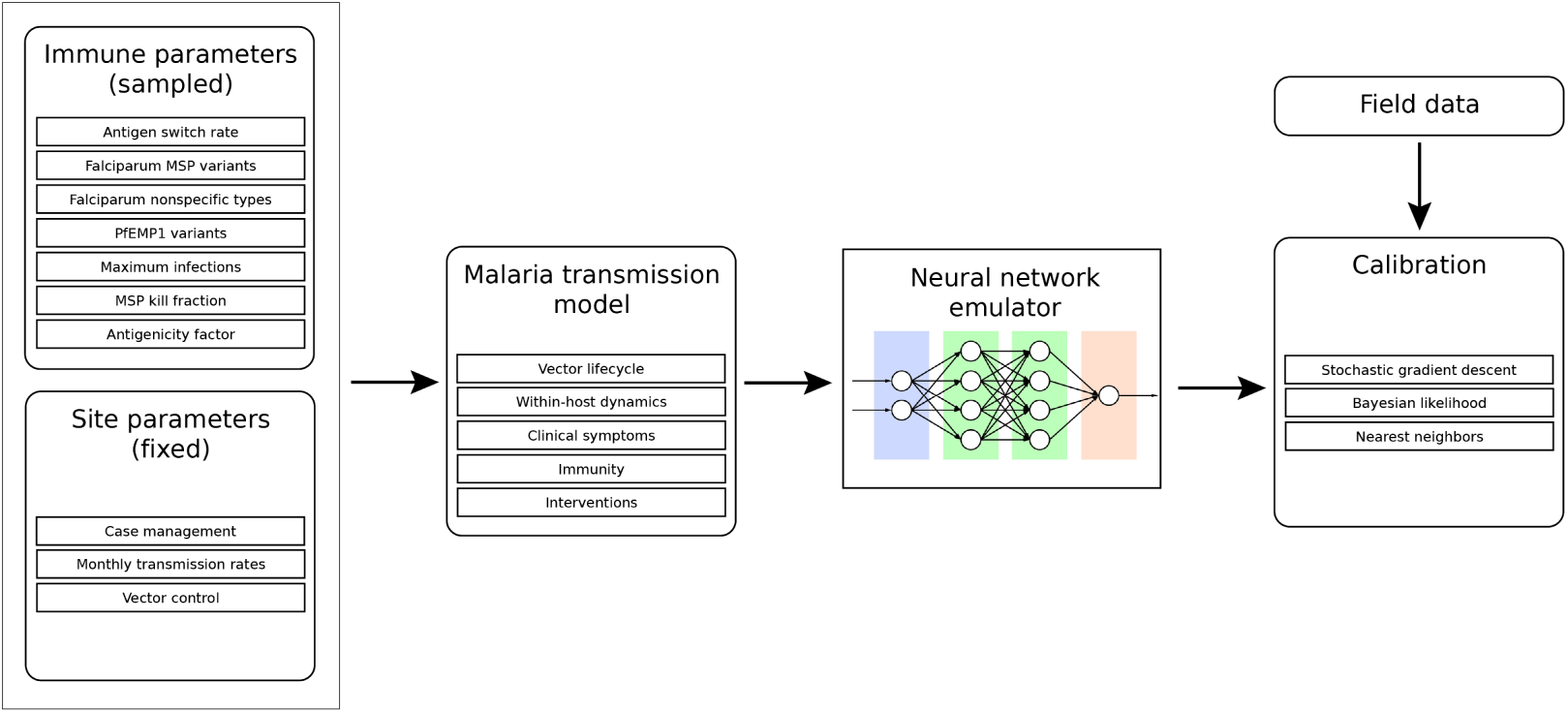
Emulation and calibration workflow. First, immune parameters are sampled, and site-specific parameters are specified. Then, the malaria transmission model (EMOD) is run for all sampled parameter sets. A neural network emulator is trained on the simulation data. Finally, calibration proceeds by comparing emulated output to field data.

#### Stochastic gradient descent

SGD allowed the emulator to explore regions of parameter space efficiently and outside the bounds on which the emulator was trained. At a high level, the algorithm proceeds as follows: 1) first, we randomly sampled 1000 initial parameter sets, 2) for each parameter set, we evaluated a loss function comparing the emulated outcomes for the parameter set against the reference, 3) we updated the parameter sets based on gradient descent (i.e., in the direction of the gradient of the loss function with respect to the parameters), 4) we iteratively continued steps 1-3 until a convergence criterion was met. Additionally, parameter values during optimization were constrained to their biologically plausible values (i.e., integer parameters are rounded, and all parameters are constrained to be positive). Finally, we selected the optimized parameter set which led to the smallest loss value. The key component of gradient descent is its iterative updating. For a parameter set **x** at iteration *t*, its update rule is given by:

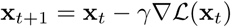

for a learning rate *γ* and loss function L. This allowed us to explore regions of parameter space that the emulator was not explicitly trained on. As before, we used the joint *ℓ*_2_ loss function:

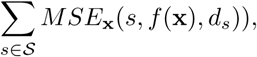

where

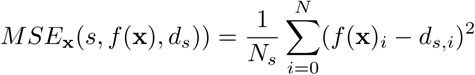

for sites *s* ∈ S, emulated outcomes *f* (**x**), and reference data for site *d_s_*. We used the Adam [34] variant of the stochastic gradient descent algorithm to efficiently search parameter space. We additionally tested different learning rates to ensure the optimization did not terminate in local extrema.

### New site inference

In addition to evaluating the emulator’s ability to capture epidemiological dynamics across study sites, we are interested in assessing i) the relationship between site-specific data and emulator performance, and ii) how applicable chosen parameter sets are to new study sites. Assessing emulator performance in the context of new field data will be important in understanding whether an emulator can capture patterns in simulation data without the need to retrain the entire system. In order to evaluate the inference ability of the emulator to unseen sites, we used a leave-one-out validation protocol, where one site is used as a held out test set. We additionally augmented the training data with site-specific epidemiological parameters such as monthly transmission rates and intervention coverages. Then, for each study site, we used the emulator not trained on that site to calibrate the remaining sites. Finally, we compared the simulation outputs corresponding to the calibrated parameter set on the site excluded during training. As models are reevaluated as new field data are obtained, this workflow will allow for an assessment of the emulator’s ability to capture dynamics in new sites, given sufficient site-specific data are included in the emulator’s training. These results are shown in the S2 Appendix.

### Sensitivity analysis

An area of concern for the developers of emulator models will be the size of the training dataset. As running complex epidemiological simulations can take long periods of time and demand large amounts of computational resources, it will be important to understand the relationship between the size of the underlying simulation suite and the performance of the emulator and calibration. To test this, we ran the emulation and calibration workflow for various subsets of the full training data and assessed the performance for each subset. Fifty replicates were run to understand the variance in performance. Our goal was to learn the smallest data subset before the emulator performance degraded. For this initial exploration, we purposefully ran a very large simulation suite, but we hope that this sensitivity analysis can shed light onto the size of simulation sets required to minimize computational resources required for future analyses. We additionally analyzed the time complexity of using an emulator compared to running the underlying mechanistic model.

## Results

### Emulator performance

First, we show the ability of the emulator to capture simulated model dynamics. After hyperparameter tuning, the final model parameters (shown in Table 3) were used to construct the neural network. Similarly, Table 4 shows the final task-specific *ℓ*_2_ losses on a holdout, previously-unseen data subset.

**Table 3.** Overview of final neural network hyperparameters.

| Parameter | Description | Value |
| --- | --- | --- |
| $n_1$ | Number of neurons in first shared layer | 512 |
| $n_2$ | Number of neurons in second shared layer | 128 |
| $n_3$ | Number of neurons in third shared layer | 4 |
| $n_4$ | Number of neurons in fourth shared layer | 64 |
| $n_5$ | Number of neurons in fifth shared layer | 16 |
| $n_6$ | Number of neurons in sixth shared layer | 32 |
| $n_7$ | Number of neurons in first task-specific layer | 16 |
| $n_8$ | Number of neurons in second task-specific layer | 2 |
| $n_9$ | Number of neurons in third task-specific layer | 8 |
| $n_{10}$ | Number of neurons in fourth task-specific layer | 256 |
| $n_{11}$ | Number of neurons in fifth task-specific layer | 8 |
| $\gamma$ | Learning rate | $1e - 4$ |
| $b$ | Batch size | 32 |
| $p$ | Dropout probability | 0.06 |
| $\lambda$ | Weight decay | $1.1e - 5$ |

**Table 4.** Overview of task-specific losses (MSE) of trained emulator.

| Task | Test set loss |
| --- | --- |
| Annual clinical incidence by age bin | 0.007 |
| Annual *P. falciparum* prevalence by age bin | 0.006 |
| Gametocytemia by age bin by month | 0.006 |
| Parasitemia by age bin by month (Matsari, Rafin Marke, Sugungum) | 0.006 |
| Parasitemia by age bin by month (Dapelogo, Laye) | 0.008 |

#### Stochastic gradient descent

Here, we show the calibrated parameter set for the stochastic gradient descent (SGD) method. This calibration methodology allows us to explore parameter sets on which the emulator was not explicitly trained. We used the SGD method to find the parameter set leading to the lowest discrepancy (loss), comparing the emulated outputs to the reference data. Then, we ran the original simulation model with the selected parameters to see the relationship between the selected parameter set, the underlying simulation, the emulated outcomes, and the reference data. Fig 2 shows the calibrated simulation, emulator, and reference data for the Dielmo, Ndiop, and Namawala study sites, corresponding to the annual incidence and prevalence outcomes, respectively. Fig 3 shows the calibrated simulation, emulator, and reference data for the Dapelogo and Laye study sites, corresponding to density and age-binned, monthly parasitemia and gametocytemia outcomes. Fig 4 shows the calibrated simulation, emulator, and reference data for the Matsari, Rafin Marke, and Sugungum study sites, corresponding to density and age-binned, monthly parasitemia outcomes. Table 5 shows the calibrated parameter set that leads to the best-fit outcomes.

**Fig 2.**
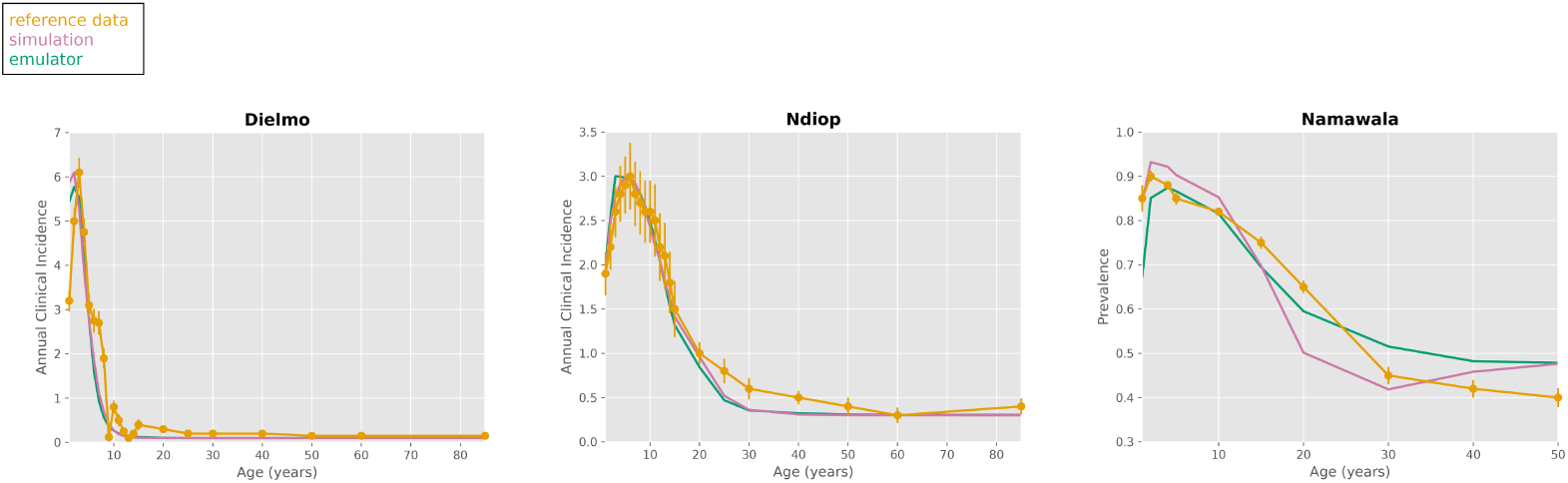
SGD calibration plots for Dielmo, Ndiop, and Namawala sites. These plots show the simulation, emulator, and reference data for the study sites corresponding to the annual clinical incidence and malaria prevalence outcomes, calibrated via the stochastic gradient descent method. The emulator was able to identify parameter sets whose outputs closely align with reference data for the sites reporting incidence data (Dielmo and Ndiop).

**Fig 3.**
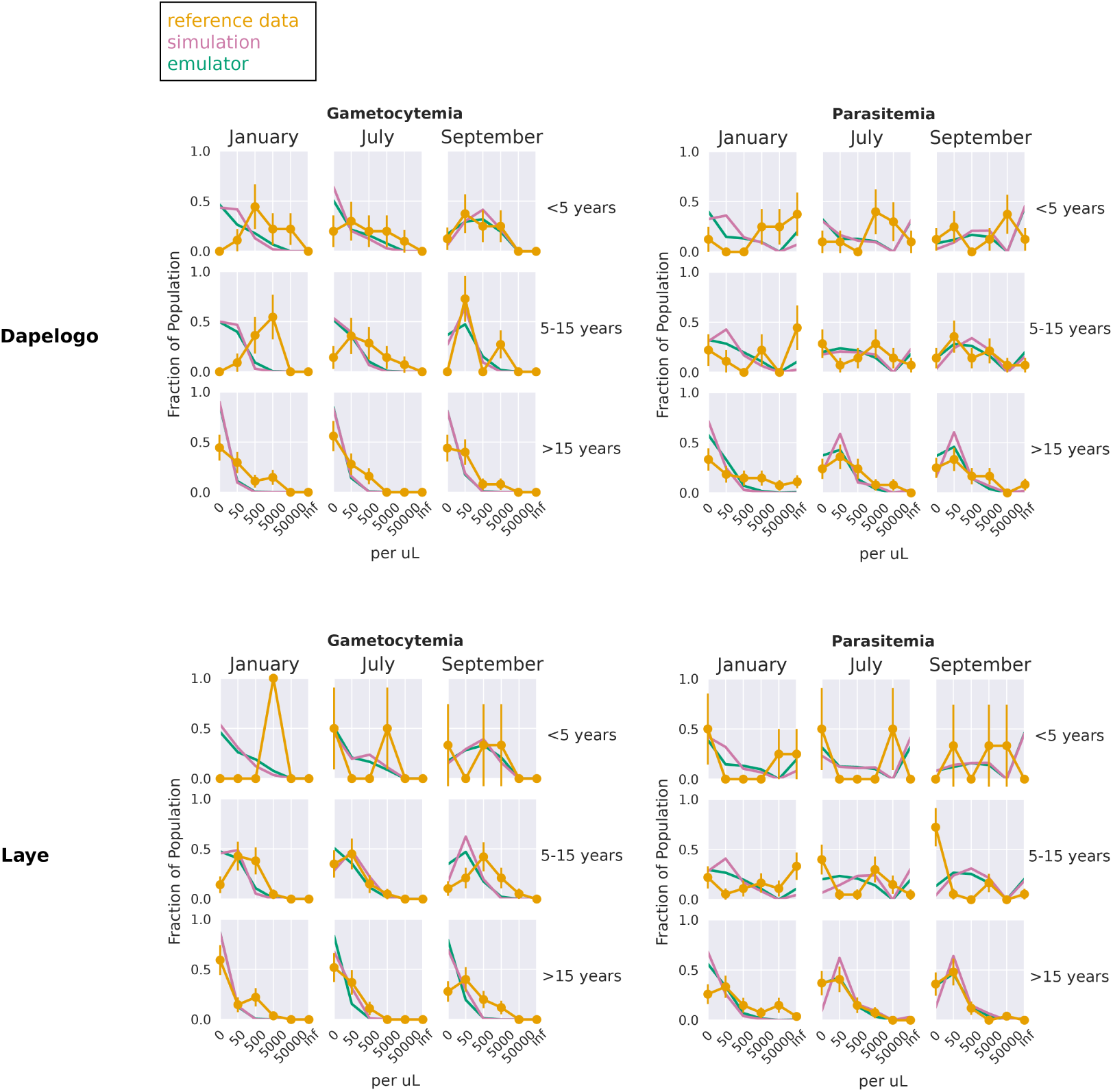
SGD calibration plots for Dapelogo and Laye sites. These plots show the simulation, emulator, and reference data for the study sites corresponding to the parasitemia and gametocytemia outcomes, calibrated via the stochastic gradient descent method.

**Fig 4.**
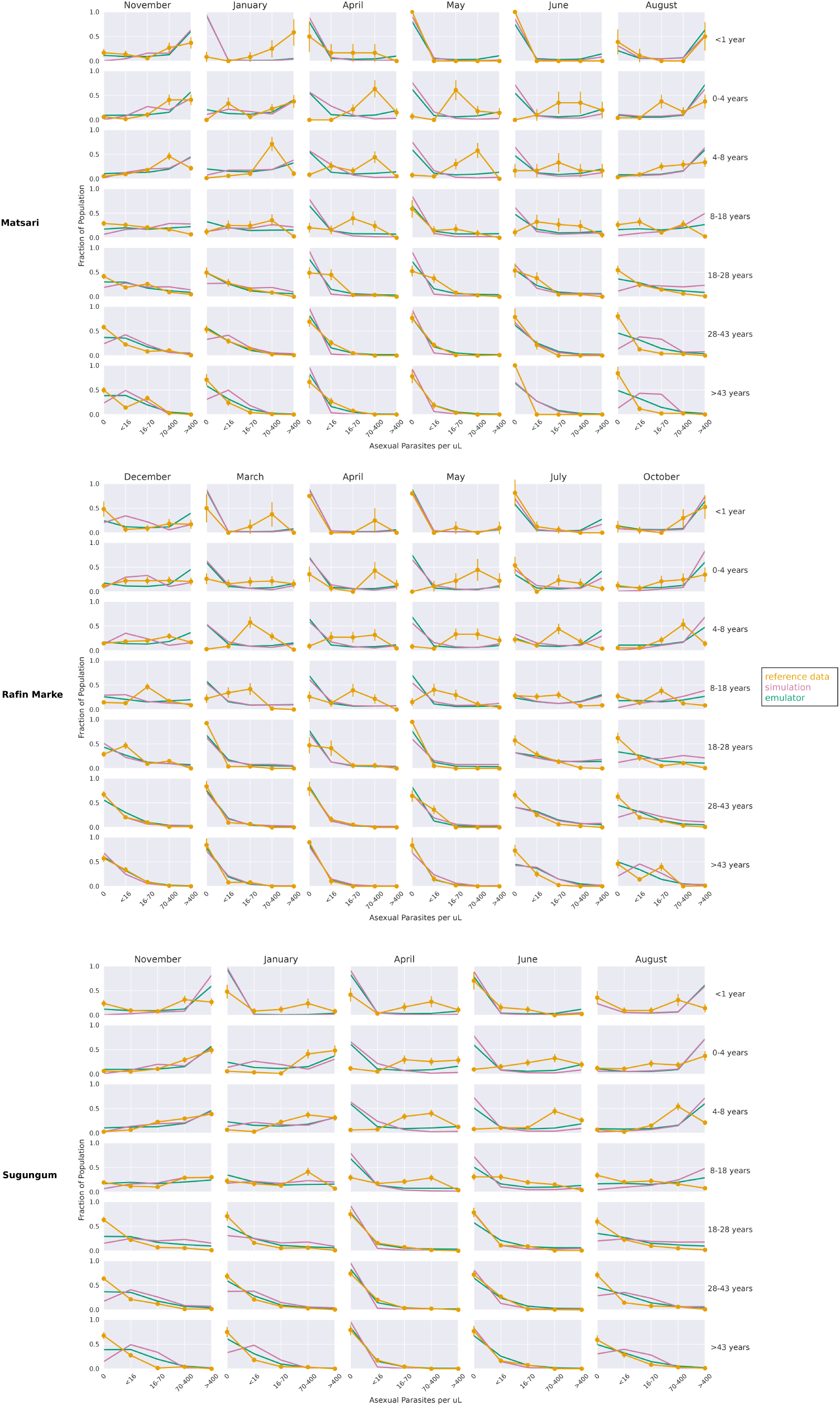
SGD calibration plots for Matsari, Rafin Marke and Sugungum sites. These plots show the simulation, emulator, and reference data for the study sites corresponding to the parasitemia outcomes, calibrated via the stochastic gradient descent method.

**Table 5.** Best fit immune parameters from SGD calibration.

| Parameter | Value |
| --- | --- |
| Antigen switch rate | $2.04e - 10$ |
| Falciparum MSP variants | 11 |
| Falciparum nonspecific types | 59 |
| Falciparum PfEMP1 variants | 1494 |
| Max individual infections | 8 |
| MSP merozoite kill fraction | 0.59 |
| Nonspecific antigenicity factor | 0.44 |

### Sensitivity analysis

Fig 5 shows the relationship between the size of the dataset on which the emulator was trained versus its ability to predict outcomes in a previously-unseen simulation data subset. Training of ML models is generally the most computationally expensive step in the inference pipeline, so there is a tradeoff between training models on more data and resource usage. For simpler models, such as the feed-forward network used to emulate EMOD, we see that the emulator can learn patterns in the simulation data even when trained on smaller data subsets; these results suggest that 40-50% of the original dataset size is sufficient to maintain similar levels of performance to the full dataset. However, for more complex models such as vision or large language models [36, 37], the size of the training data is more relevant. Additionally, we see that providing the model with too much training data leads to an increase in the variance of loss for the unseen data subset, which is likely due to the smaller test set or overfitting.

**Fig 5.**
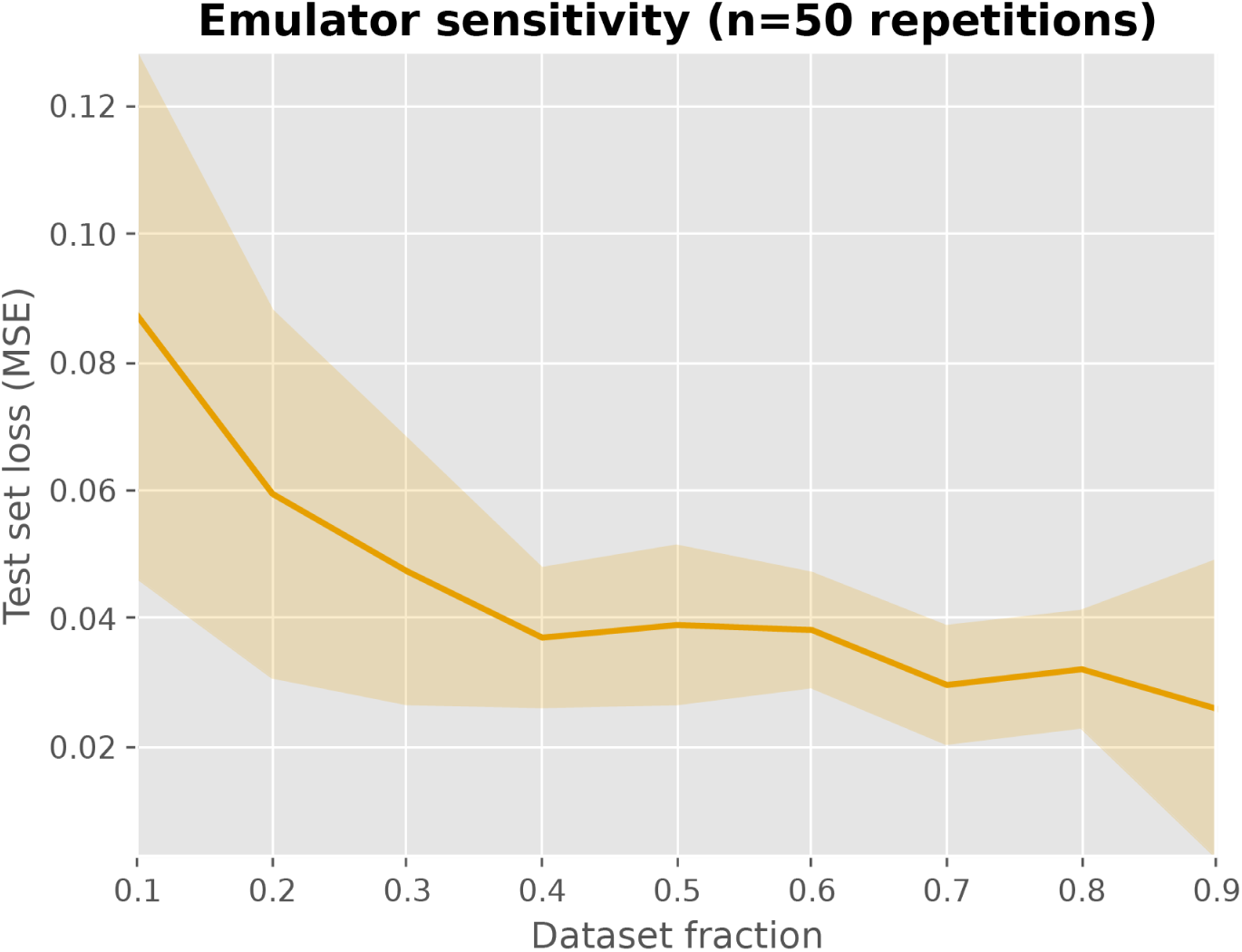
Sensitivity analysis for emulator training dataset size. 50 replicates were run to evaluate the relationship between the training data size and the emulator performance on a hold-out test set.

### Time complexity analysis

A primary motivation of employing emulator models is their relative computational efficiency [38]. Once a neural network is trained, inference (i.e., obtaining outputs from a set of inputs) is generally orders of magnitude faster than running the underlying simulation. Training and selecting hyperparameters of DNNs generally are the rate-limiting steps in an ML pipeline; in our case, a full hyperparameter search took roughly eight hours on a single graphics processing unit (GPU). However, advances in data parallelization [39] can significantly reduce training times for DNNs and other network architectures. To demonstrate the inference speed of the emulator as compared to the underlying simulations, we show the distribution of runtimes of the same 2000 parameter sets, segmented by study site in Fig 6. We see that the trained emulator can, for the same parameter sets, predict outcomes several orders of magnitude faster than the runtimes of the underlying simulations. Additionally, simulation runtimes show more variance within and across study site, as they are highly sensitive to input parameters. For EMOD, parameters such as the maximum number of infections and parasite variants can significantly expand the runtime of a simulation, as more infection scenarios need to be considered explicitly. A trained emulator conversely shows little variance and can efficiently handle the same parameter sets.

**Fig 6.**
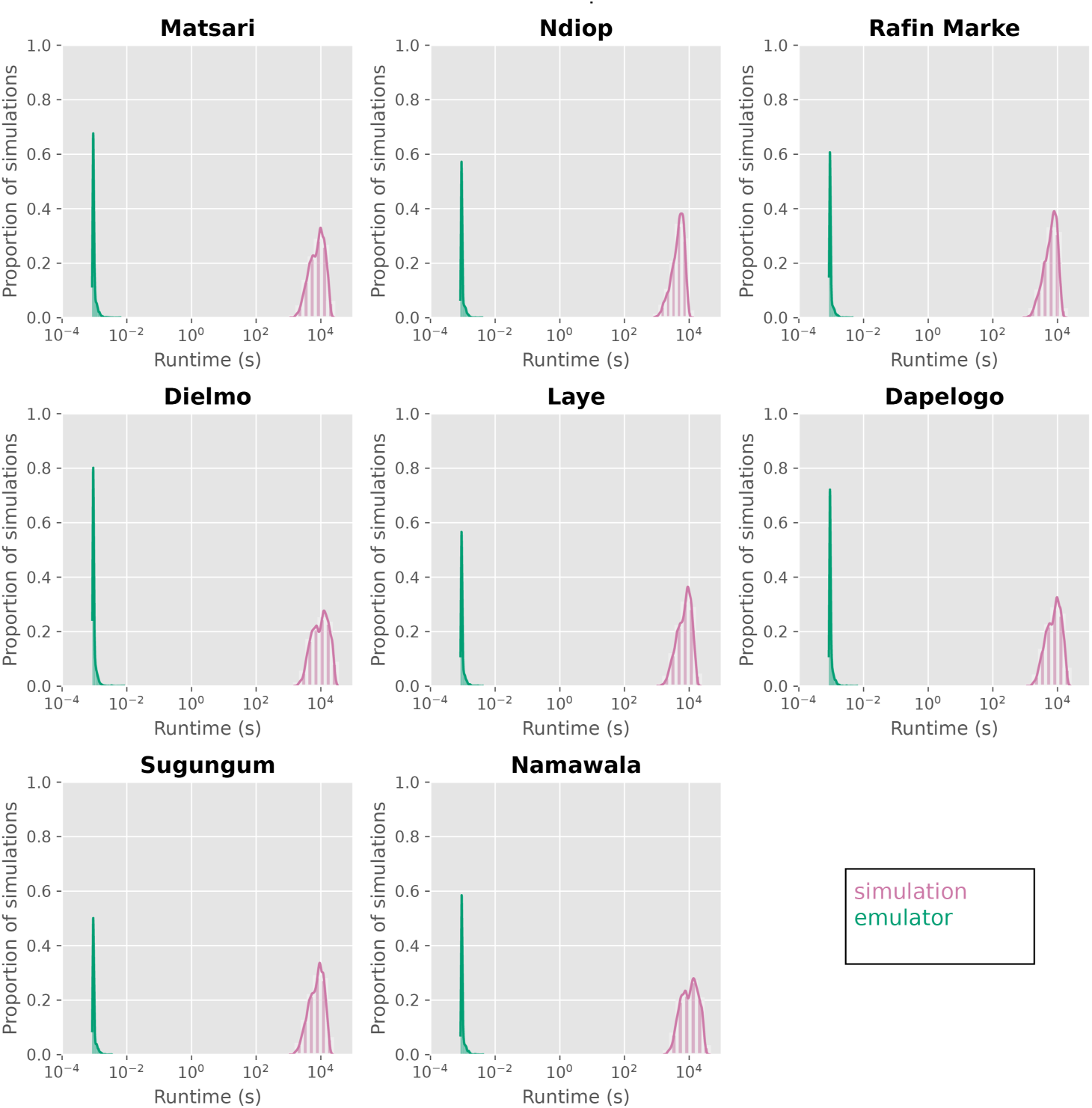
Comparison of runtimes between emulated and simulated outcomes, segmented by site. Trained emulators can predict outcomes several orders of magnitude faster than the runtimes of the underlying simulation.

## Discussion and future directions

Here, we have described a workflow to 1) emulate a complex malaria transmission model using deep neural networks, and 2) calibrate the underlying mechanistic model to reference data using the emulator. Supplementary work showed the ability of DNN emulators to infer outputs to study sites not seen during training. Through careful sampling of the underlying parameter space, calibration methodology, and neural network architecture, we have shown that machine learning alternatives to traditional model calibration techniques (e.g., Markov chain Monte Carlo, particle filters) can be used to rapidly search parameter space and align mechanistic models to field data.

There are a couple of issues that may make calibration more challenging when using an emulator for calibration. Specifically, an emulator may smooth over some detailed model dynamics, leading to potential biases or reduced accuracy in parameter estimates, particularly for parameters with subtle or complex effects. Additionally, the emulator fitting itself introduces another layer of uncertainty, which could propagate into the calibration process. However, despite these challenges, emulator-based calibration remains highly beneficial and robust. Emulators significantly reduce computational time, allowing exploration of a broader parameter space and enabling more extensive uncertainty quantification than would be possible with direct simulations alone.

Furthermore, with careful validation - such as we conducted in this work - the emulator closely approximates the full model, preserving essential dynamics while enabling efficient calibration.

In terms of model architecture, we employed a multitask, feed-forward neural network mapping the input parameters (in this case, parameters associated with immunological dynamics of malaria transmission) to age and density-binned epidemiological outcomes. While more complex architectures exist (e.g., recurrent or transformer-based models) for time series inference, these models require more resources to train properly. In designing the emulator, we opted to employ a multitask strategy (i.e., one emulator jointly predicts all outcomes) for two primary reasons. First, this paradigm allows the emulator to learn shared patterns across the input and output space, as each outcome is likely not biologically independent of the others. In addition, model complexity and computational resource requirements are lower in training a single, multitask emulator as compared to several individual emulators.

We identified two primary challenges in designing a multitask emulator that model developers should consider when applying this framework to their simulation data: task weighting and specification of the output data format. With respect to task weighting, we opted to weight each task equally, as to not introduce additional complexity into the training pipeline. Prior research has found that model performance can be highly sensitive to model weights [40], and the weights themselves can be treated as a tuneable parameter. Several strategies have been proposed to address task weighting to achieve optimal network performance [24, 40]. While we were able to achieve adequate performance with a uniform loss weighting scheme, the loss weights are nonetheless another set of hyperparameters that should be considered when designing an emulator training pipeline. Specifying the output data format posed another challenge: we deliberately formatted the emulator’s output to match the dimensionality of available field data. As new data become available, however, a key concern is the usability of the emulator with data formats that differ from those on which the network was trained. Recent developments in transfer learning and model finetuning [41] provide a framework by which the underlying emulator can still be used to align to new data without the need to retrain the entire system, via updating the neural network’s output layers. These techniques could be leveraged here to align the emulator with new field data without needing to rerun the training pipeline.

We have additionally shown that neural network emulator models can predict outcomes for previously-unseen sites, allowing for more flexibility as more field data are obtained. Though the emulator performance is slightly degraded due to less training data and extrapolation to new study sites, the fits suggest that the emulator can still narrow down parameter space to reasonable values, given that the emulator training is augmented with suitable site-specific data such as transmission rates and intervention coverages. In our analysis, the structure of the field data matches the simulation data on which the emulator was trained, but future directions could consider updating the final layers of the neural network to output new data structures, as is done in other domains [41, 42]. This would allow even more flexibility when calibrating new field sites.

A concern with the usability of an emulator-based calibration workflow regards its ability to infer outcomes in parameter spaces not seen during emulator training. While we have shown that SGD allows for reasonable inference on parameter sets not explicitly seen during training, this method generally stays within the bounds of the training set, essentially interpolating between sampled points. To evaluate generalizability even further, we conducted supplementary analyses (S2 Appendix) to test the emulator’s ability to calibrate data from study sites not seen during training. In this framework, an emulator was trained on simulation data from all study sites, excluding one. Then all sites were calibrated using the emulator. Our preliminary results show that the emulator is still able to capture relationships across all study sites, but its performance is degraded compared to emulators trained with the full dataset. Future iterations could examine training on broader site characteristics (e.g., variable EIRs, intervention coverages, etc.) to improve generalizability. Additionally, the strength of DNN-based emulators lies in their ability to leverage hierarchical learning, capturing nonlinear parameter interactions through layers of progressively abstracted features. With respect to sampling the parameter space, Rather than explicitly sampling every possible region in the parameter space uniformly, DNNs identify and generalize patterns across parameter sets, allowing them to effectively interpolate and extrapolate from limited training data [43]. This capability enables DNNs to efficiently represent high-dimensional spaces without exhaustive sampling, making them particularly suitable for scaling up from a moderate number of parameters to a higher-dimensional range.

With respect to the emulator’s performance and the size of the training data, our study found that smaller parameter sets can still capture the underlying simulation’s dynamics, albeit with more variance. This can inform the design of emulators in the context of limited computational resources.

### Interpretability of deep neural network models

A major challenge with the acceptance of DNN models is their interpretability [44]. That is, most traditional DNNs are trained by minimizing the loss between the training data and the predicted outputs; the mechanisms on *how* they learn these patterns can be hard to discern, especially as model complexity increases. For some applications, understanding the underlying mechanisms of a DNN may be less of a concern. But for epidemiological models, the mechanisms of disease transmission are important to understanding how interventions are expected to behave on outcomes of interest.

Additionally, in the calibration phase, it is important to ensure that model inputs and outputs correspond to biologically plausible parameter values (for example, by constraining values to realistic ranges, even if other values improve numerical performance).

An active area of machine learning research is improving the interpretability of models. Methods include reinforcement learning with human feedback and mixture of experts, among others [45, 46]. In the disease modeling realm, trusting inferences from ML models will require an understanding of the mechanisms by which an inference is made. Fine-tuning may also be required to ensure the model outputs are aligned with biologically plausible values. In our work, we have shown that ML emulators can rapidly search parameter space in order to align model outputs with reference data.

Calibration outputs should be assessed against domain knowledge to identify which parameter sets are the most feasible given transmission setting and model dynamics [18]. Recent methods have been introduced to even incorporate domain knowledge into the training of ML models [47].

### Comparison of calibration techniques

Here we have shown coupling SGD with a trained emulator is able to produce reasonable calibration fits across sites and outputs, comparing emulated outputs to reference data. In supplementary analyses (S1 Appendix), we evaluated two additional parameter selection techniques, involving Bayesian likelihoods and a nearest neighbors approach. In choosing a calibration method, attention must be paid to the functional form (if any) imposed on the outputs and how the method traverses parameter space. In the case of Bayesian likelihood and nearest neighbors, parameter sets used to train the emulator were sampled, and their associated outputs were compared against the reference data. Bayesian inference additionally requires a functional form of the conjugate distribution, which we derived from prior work [25]. Both these methods are rapid to evaluate, as they require a simple likelihood or loss calculation in comparison to reference data. For this analysis, Bayesian likelihoods were calculated on a previously-sampled parameter set. It could also be coupled with SGD as a “loss function” to provide the flexibility of evaluating new parameter sets while imposing a functional form on the data. Stochastic gradient descent with an MSE loss function, conversely, does not specify any functional form of the outputs nor does it require a sampling of the parameter space beforehand. Thus, SGD and other numerical optimization methods may be more flexible and result in better calibration fits, particularly those that are out-of-distribution (i.e., outside the emulator’s training manifold). Numerical optimization methods do however require additional attention to hyperparameters, as the choice of initial values, learning rates, and momentum can greatly impact final fits [48]. Nonetheless, these methods serve as powerful and flexible tools in emulator-based calibration problems. Additional methods not evaluated here, such as iterative adaptive sampling [49], have shown promise in multitask calibration of agent based models.

### Discussion of emulation frameworks

Here, we provide a brief overview of two main surrogate modeling techniques employed in infectious disease analysis and compare them to the DNN-based approach utilized in this study. Gaussian processes (GPs) represent a stochastic process (generally indexed by time or space), whose finite subsets follow a multivariate normal distribution. They tend to have fewer hyperparameters compared to other methods (and thus are easier to train) and can be used to model a wide range of scientific processes. Additionally, calibration via GPs has been shown to be robust to convergence issues and GPs provide a natural quantification of model uncertainty [50, 51]. Neural network-based approaches, on the other hand, can be more flexible in that they do not impose a normality assumption on the training data, but do require (often orders of magnitude) more parameters and attention to hyperparameters during training. Recent surrogate modeling literature has introduced Bayesian neural networks as a method to simultaneously model the solution space and uncertainty, similar to GPs [18, 30, 31]. We opted for a traditional DNN-based approach to demonstrate its flexibility to highly non-linear simulation data and rapid inference. GP-based emulation studies [50] have additionally employed adaptive sampling to further refine parameter space and quantify uncertainty. DNN emulators can integrate effectively with adaptive sampling frameworks, employing uncertainty quantification methods such as ensembles or Monte Carlo Dropout [52, 53]. Hence, the potential for adaptive sampling is not only maintained but potentially enhanced by a DNN approach, given the computational efficiency and scalability inherent to these methods. Recent research [54, 55] has demonstrated that deep neural networks (DNNs) combined with effective methods of uncertainty estimation such as the Δ-UQ approach or Direct Epistemic Uncertainty Prediction (DEUP) often yield better performance in adaptive sampling and Bayesian optimization tasks compared to traditional GPs, Bayesian neural networks, and deep ensembles. In future work, we aim to systematically benchmark our neural-network emulator against alternative surrogate models—carefully exploring and tuning each model’s hyperparameter space to ensure a fair, rigorous comparison to the same reference dataset.

### Limitations

This study has several limitations. Mechanistic models of malaria transmission must be sufficiently complex to capture the seasonal and heterogeneous (by age, species, and location) nature of the disease. Even with complex models, capturing age-based heterogeneity is difficult [1] due to the nature of immune dynamics in children (i.e., children often experience more severe cases of malaria after maternal immunity has waned, but exposure immunity has not yet developed). Our calibration methodology aimed to find one set of immune parameters that best fit the reference data across age groups, but we see that the fits for parasite density are markedly worse in juvenile age groups, particularly for the Garki sites. This finding suggests that one set of immune parameters may not be fully appropriate across the age spectrum and that juvenile dynamics should be calibrated separately. Additionally, the underlying mechanistic model may not be fully describing immune dynamics in these age strata. Both of these aspects limit the ability of the calibration workflow to extend to younger age groups. From an intervention planning perspective, these dynamics should be better understood, as interventions often target children to reduce the significant morbidity and mortality they face from malaria.

A central challenge concerns the principled development of DNNs and scientific process emulators. Many architectures, hyperparameters, and training data sampling schemes must be considered. While some tools exist to help design these models, many choices must be made beforehand to ensure the emulator is capturing the dynamics of the transmission model. For other use cases where the mechanisms of the DNN are important to understand, careful attention must be paid to the development of the emulator to ensure it accurately represents the underlying scientific process [18].

Additionally, the tradeoff between simulation dataset size and model accuracy must be considered in the context of limited computational resources. We deliberately sampled a large range of parameter space as a proof-of-concept, but future studies should explore more deeply the ability of the emulator to learn simulation dynamics with more limited training datasets. In generating the training dataset for the primary analysis, we assumed that site characteristics (e.g. monthly EIRs, intervention coverages, etc.) were static across simulations. While this assumption may not be valid under all circumstances, such as interventions that may influence EIR via decreased prevalence, field data measuring EIR can be challenging to collect and validate. Our supplementary analyses S2 Appendix show that augmenting the input data with site-specific EIRs can allow for calibration to new sites. In this transfer learning paradigm, the emulator learns relationships between site-specific characteristics like EIR and simulation outcomes. Pre-trained emulators can then be used to infer outcomes on new sites, given appropriate EIR data are supplied, without the need to retrain the emulator. Our preliminary analyses show that this approach can allow for refinement of calibration parameter space for sites not seen during training and incorporation of site-specific metadata to improve parameter estimates.

A major computational tradeoff in designing an emulator-based calibration workflow regards the generation of a robust training dataset. Fig 6 shows that once an emulator is trained, inference can be conducted rapidly. However, the training process itself requires two computationally-heavy steps: i) generating a suite of simulations and, ii) identifying optimal emulator hyperparameters. Deep learning techniques, such as fine-tuning and transfer learning, substantially mitigate issues of long training times. Extensive literature demonstrates that fine-tuning pre-trained networks can significantly reduce computational demands and improve training efficiency without sacrificing inference quality [56, 57]. Moreover, our current implementation utilized a single GPU; employing more advanced GPUs or multiple GPUs in parallel could substantially decrease training duration and further enhance computational efficiency. One example is the article by Goyal *et al.* [58], where they showed how convolutional neural networks (CNNs), a natural extension of the MLP architecture we use in this work, can essentially train large volumes of image data in an hour. Our initial experiments already suggest promising outcomes with such approaches.

A final concern relates to the validation of the calibration workflow. Biological parameters associated with malaria transmission can be difficult to quantify. Drawing from previous literature [25, 60], conservative bounds were chosen to fully capture the parameter space associated with each biological parameter. As such, simulation times varied significantly, as wider parameter ranges led to a larger variation in simulation scenarios (Fig 6 shows the distribution of simulation runtimes). With refined estimates of these parameters, either from literature or biological experiments, tighter LHS ranges could be considered for future iterations, as both simulation and calibration efficiency would increase significantly. DNN emulators can offer a very favorable trade-off between computational efficiency and reliable predictions, thereby ensuring accurate and policy-relevant calibration outcomes. Moreover, transfer learning is a technique by which neural networks could outperform other competing surrogates, where one can potentially bootstrap multiple related datasets (e.g., simulations and field data) or even simulations of varying fidelities (i.e., data structures) to effectively predict on the dataset of interest. This would not only greatly reduce the amount of data required, but also provide a meaningful way to capture implicit characteristics in datasets into the calibration process. There are numerous successful examples in literature in diverse applications. In malaria for example, transfer learning has been used to train DNNs for imaging and data analysis [61, 62], which makes a case for DNN-based calibration taking advantage of transfer learning.

## Supporting information

**S1 Appendix. Nearest neighbors and likelihood calibrations for all sites**. Calibration results for complete (all-site) dataset.

**S2 Appendix. Site-excluded calibrations**. Calibration results for inferring outcomes on previously-unseen sites in emulator training.

## Code availability statement

Model weights, training, and calibration code are available on GitHub.

