## Supplementary material for "Multitask deep learning for the emulation and calibration of an agent-based malaria transmission model": S1 Appendix

### S1 Appendix - Likelihood and nearest neighbors calibration

#### Calibration

The following plots show the calibration results for the likelihood and nearest neighbors approaches for the complete (i.e., all-sites) dataset.

##### Bayesian likelihood maximization

We used a Bayesian likelihood-based approach to find the parameter set which maximized the joint likelihood across sites as compared to the reference data. This approach, previously detailed [1–3], is described briefly here. For each data type, an initial uniform prior is updated by the simulated (or emulated) outcomes. Then, a likelihood is calculated against the reference data by marginalizing over the true parameter of the prior distribution. Incidence is modeled as a gamma-Poisson conjugate distribution, prevalence is modeled as a beta-binomial distribution, and parasite/gametocyte density is modeled as a Dirichlet-multinomial distribution. The joint likelihood is the product of likelihoods for each age group, month, study site, and parasite bin, which is also multiplied with the likelihoods from prevalence and incidence outcomes to obtain the total likelihood of the parameter set. The selected parameter set maximizes the joint likelihood across sites.

###### Model fits

We show the calibrated parameter set for the likelihood-based approach to aligning simulation data with reference data. Fig 1 shows the calibrated simulation, emulator, and reference data for the Dielmo, Ndiop, and Namawala study sites, corresponding to the annual incidence and prevalence outcomes, respectively. Fig 2 shows the calibrated simulation, emulator, and reference data for the Dapelogo and Laye study sites, corresponding to density and age-binned, monthly parasitemia and gametocytemia outcomes. Fig 3 shows the calibrated simulation, emulator, and reference data for the Matsari, Rafin Marke, and Sugungum study sites, corresponding to density and age-binned, monthly parasitemia outcomes. Table 1 shows the calibrated parameter set giving rise to the best-fit outcomes.

| **Parameter** | **Value** |
|---|---|
| Antigen switch rate | 0.18 |
| Falciparum MSP variants | 154 |
| Falciparum nonspecific types | 305 |
| Falciparum PfEMP1 variants | 3651 |
| Max individual infections | 9 |
| MSP merozoite kill fraction | 0.83 |
| Nonspecific antigenicity factor | 0.46 |

**Table 1. Best fit immune parameters from likelihood calibration.**

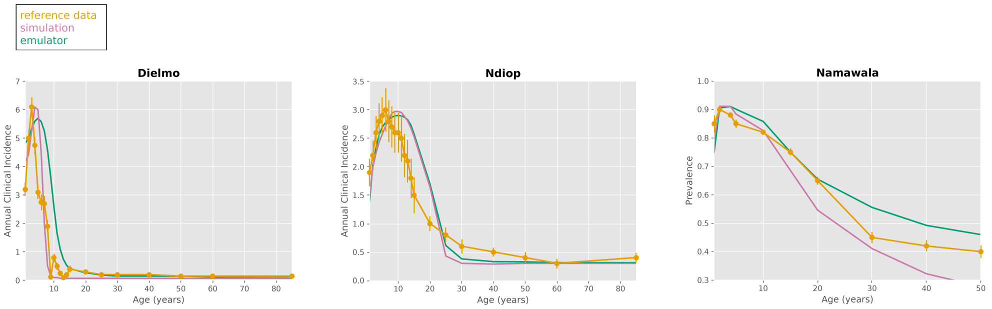


**Fig 1. Likelihood calibration plots for Dielmo, Ndiop, and Namawala sites.** These plots show the simulation, emulator, and reference data for the study sites corresponding to the annual clinical incidence and malaria prevalence outcomes, calibrated via the likelihood method.

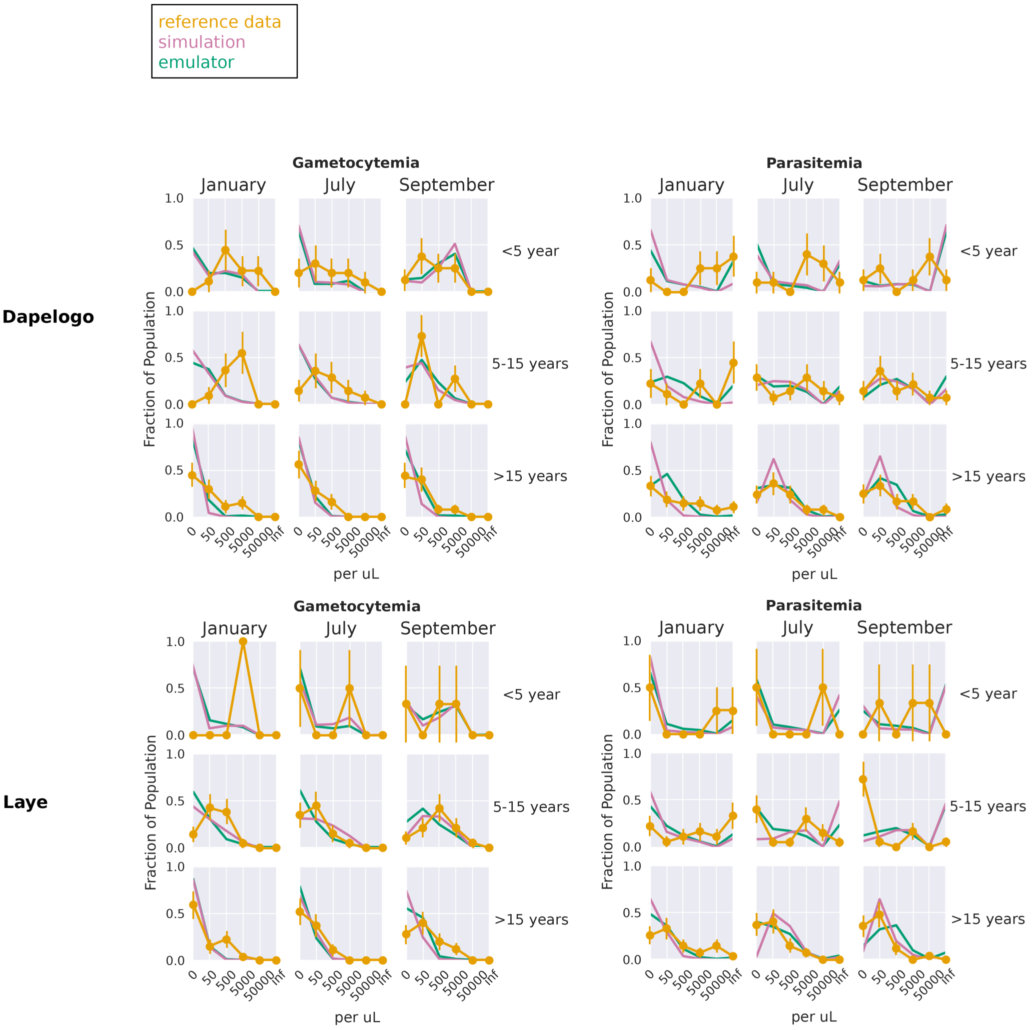


**Fig 2. Likelihood calibration plots for Dapelogo and Laye sites.** These plots show the simulation, emulator, and reference data for the study sites corresponding to the parasitemia and gametocytemia outcomes, calibrated via the likelihood method.

#### Nearest neighbors

We additionally tested finding the parameter set which minimized the $\ell_2$ distance between emulated outcomes and the reference data across sites, in order to find the parameter set whose emulated outcomes are nearest to the reference data. As stated previously, the $\ell_2$ distance is the represents the mean squared numerical discrepancy between two vectors. Here, the goal is to minimize the total $\ell_2$ distance given by:

$$\min_{\mathbf{x}} \sum_{s \in \mathcal{S}} MSE_{\mathbf{x}}(s, f(\mathbf{x}), d_s)),$$

where

$$MSE_{\mathbf{x}}(s, f(\mathbf{x}), d_s)) = \frac{1}{N_s} \sum_{i=0}^{N} (f(\mathbf{x})_i - d_{s,i})^2$$

for sites $s \in \mathcal{S}$, emulated outcomes $f(\mathbf{x})$, and reference data for site $d_s$.

##### Model fits

Here we show the parameter set whose emulated outputs were nearest to the reference data with respect to the $\ell_2$ loss criterion. Fig 4 shows the calibrated simulation, emulator, and reference data for the Dielmo, Ndiop, and Namawala study sites, corresponding to the annual incidence and prevalence outcomes, respectively. Fig 5 shows the calibrated simulation, emulator, and reference data for the Dapelogo and Laye study sites, corresponding to density and age-binned, monthly parasitemia and gametocytemia outcomes. Fig 6 shows the calibrated simulation, emulator, and reference data for the Matsari, Rafin Marke, and Sugungum study sites, corresponding to density and age-binned, monthly parasitemia outcomes. Table 2 shows the calibrated parameter set giving rise to the best-fit outcomes.

| **Parameter** | **Value** |
|---|---|
| Antigen switch rate | $4.6e-06$ |
| Falciparum MSP variants | 6 |
| Falciparum nonspecific types | 300 |
| Falciparum PfEMP1 variants | 1739 |
| Max individual infections | 9 |
| MSP merozoite kill fraction | 0.67 |
| Nonspecific antigenicity factor | $9.4e-06$ |

**Table 2. Best fit immune parameters from nearest neighbors calibration.**

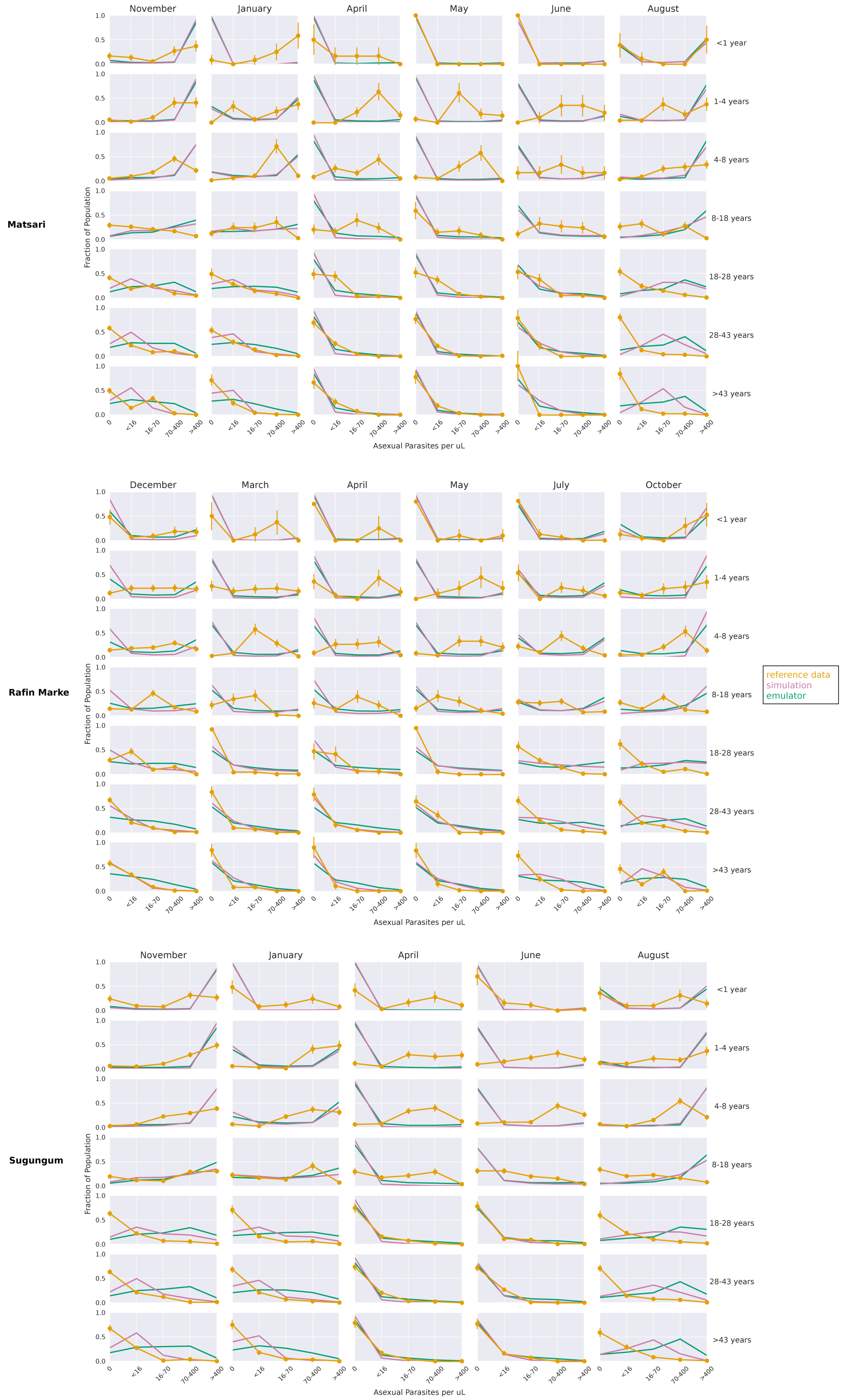


**Fig 3. Likelihood calibration plots for Matsari, Rafin Marke and Sugungum sites.** These plots show the simulation, emulator, and reference data for the study sites corresponding to the parasitemia outcomes, calibrated via the likelihood method.

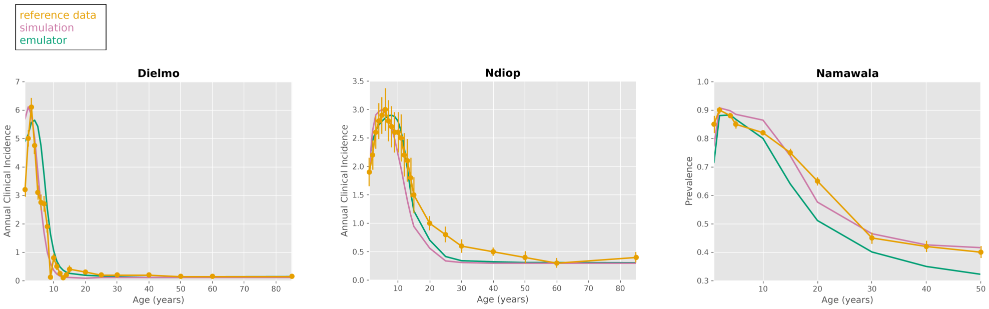


**Fig 4. Nearest neighbor calibration plots for Dielmo, Ndiop, and Namawala sites.** These plots show the simulation, emulator, and reference data for the study sites corresponding to the annual clinical incidence and malaria prevalence outcomes, calibrated via the nearest neighbors method.

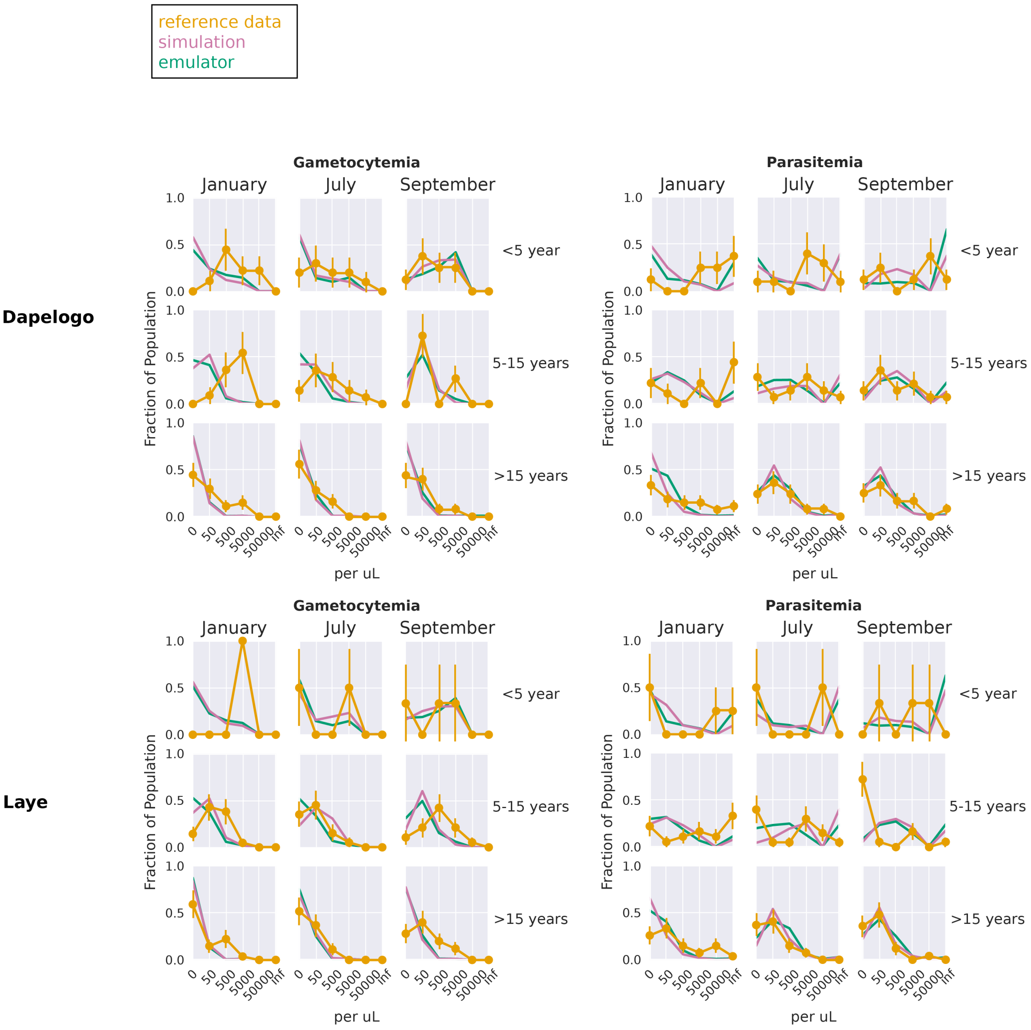


**Fig 5. Nearest neighbor calibration plots for Dapelogo and Laye sites.** These plots show the simulation, emulator, and reference data for the study sites corresponding to the parasitemia and gametocytemia outcomes, calibrated via the nearest neighbors method.

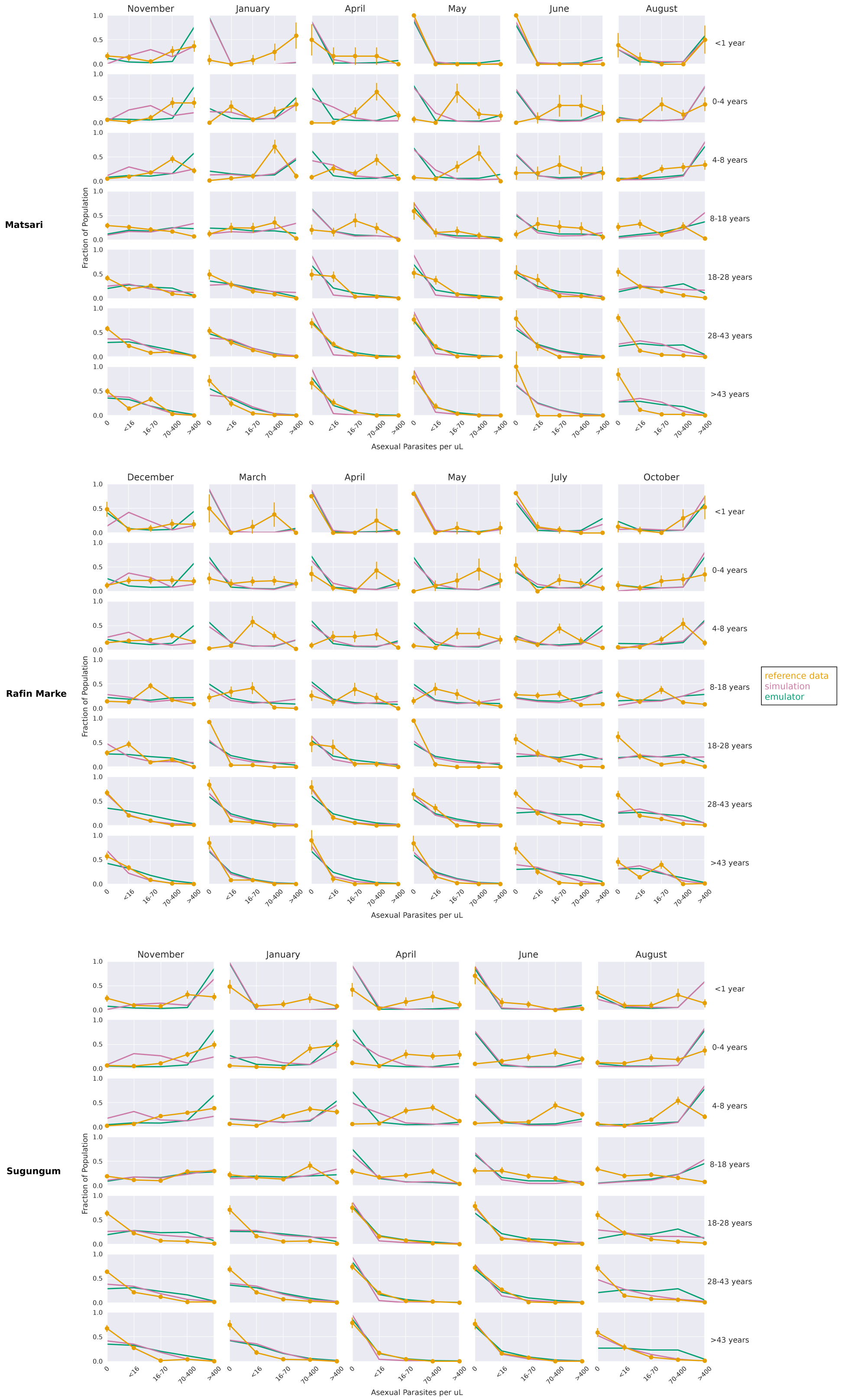


**Fig 6. Nearest neighbor calibration plots for Matsari, Rafin Marke and Sugungum sites.** These plots show the simulation, emulator, and reference data for the study sites corresponding to the parasitemia outcomes, calibrated via the nearest neighbors method.
