## Supplementary material for "Multitask deep learning for the emulation and calibration of an agent-based malaria transmission model": S2 Appendix

### S2 Appendix - Site-excluded calibration

#### New site inference

For each study site, an emulator was trained on all simulation data, excluding the data for the site. Inputs were augmented with site-specific data such as monthly transmission rates and interventions. Calibration were conducted across all sites (excluding the new site), and the underlying simulation was run with the calibrated parameters for all sites. The subsequent figures show results for stochastic gradient descent calibrations. Fig 1 shows the calibrated simulation, emulator, and reference data for the Dielmo, Ndiop, and Namawala study sites, corresponding to the annual incidence and prevalence outcomes, respectively. Fig 2 shows the calibrated simulation, emulator, and reference data for the Dapelogo and Laye study sites, corresponding to density and age-binned, monthly parasitemia and gametocytemia outcomes. Fig 3 shows the calibrated simulation, emulator, and reference data for the Matsari, Rafin Marke, and Sugungum study sites, corresponding to density and age-binned, monthly parasitemia outcomes.

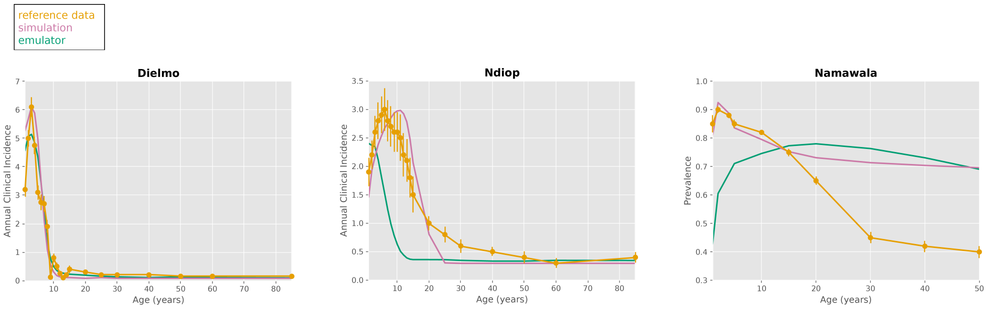


**Fig 1. Site-excluded SGD calibration plots for Dielmo, Ndiop, and Namawala sites.** These plots show the simulation, emulator, and reference data for the study sites corresponding to the annual clinical incidence and malaria prevalence outcomes, calibrated via the stochastic gradient descent method, using emulators not explicitly trained on data from these sites.

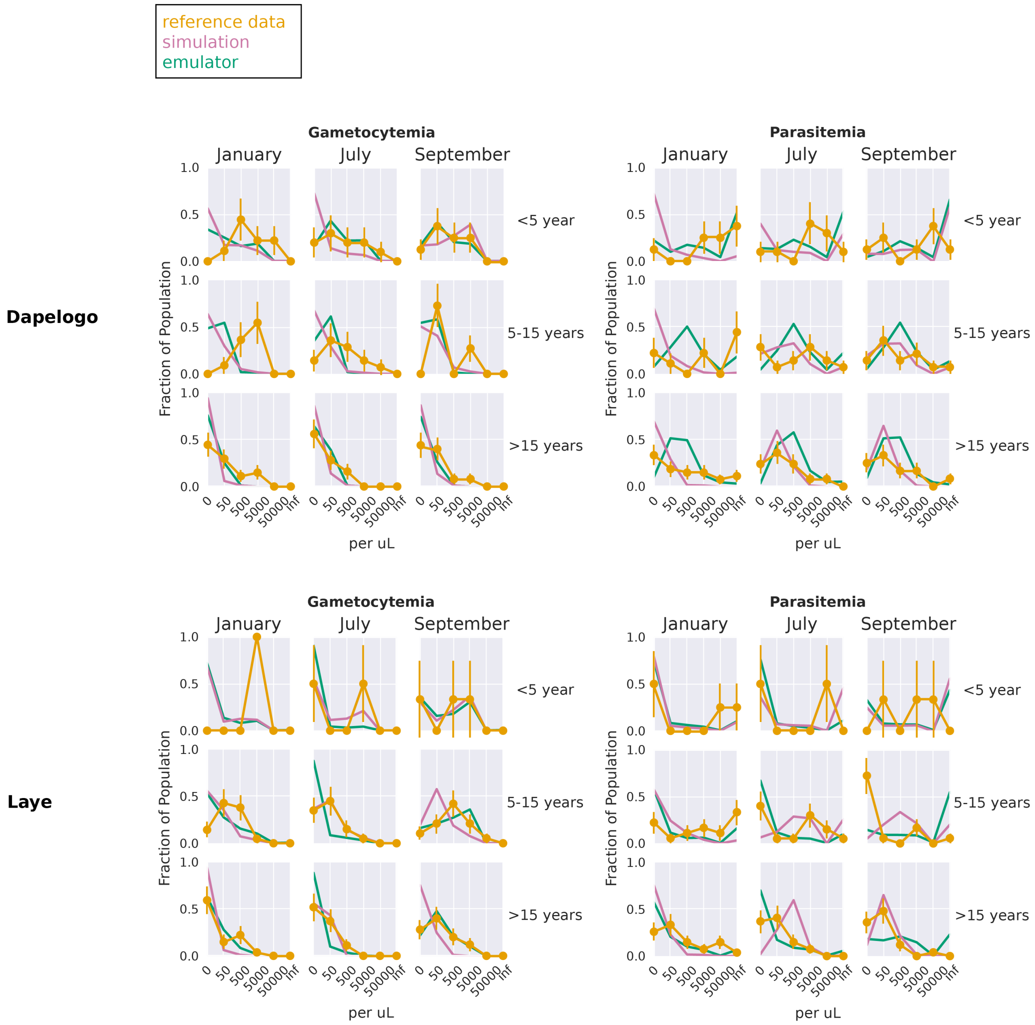


**Fig 2. Site-excluded SGD calibration plots for Dapelogo and Laye sites.** These plots show the simulation, emulator, and reference data for the study sites corresponding to the parasitemia and gametocytemia outcomes, calibrated via the stochastic gradient descent method, using emulators not explicitly trained on data from these sites.

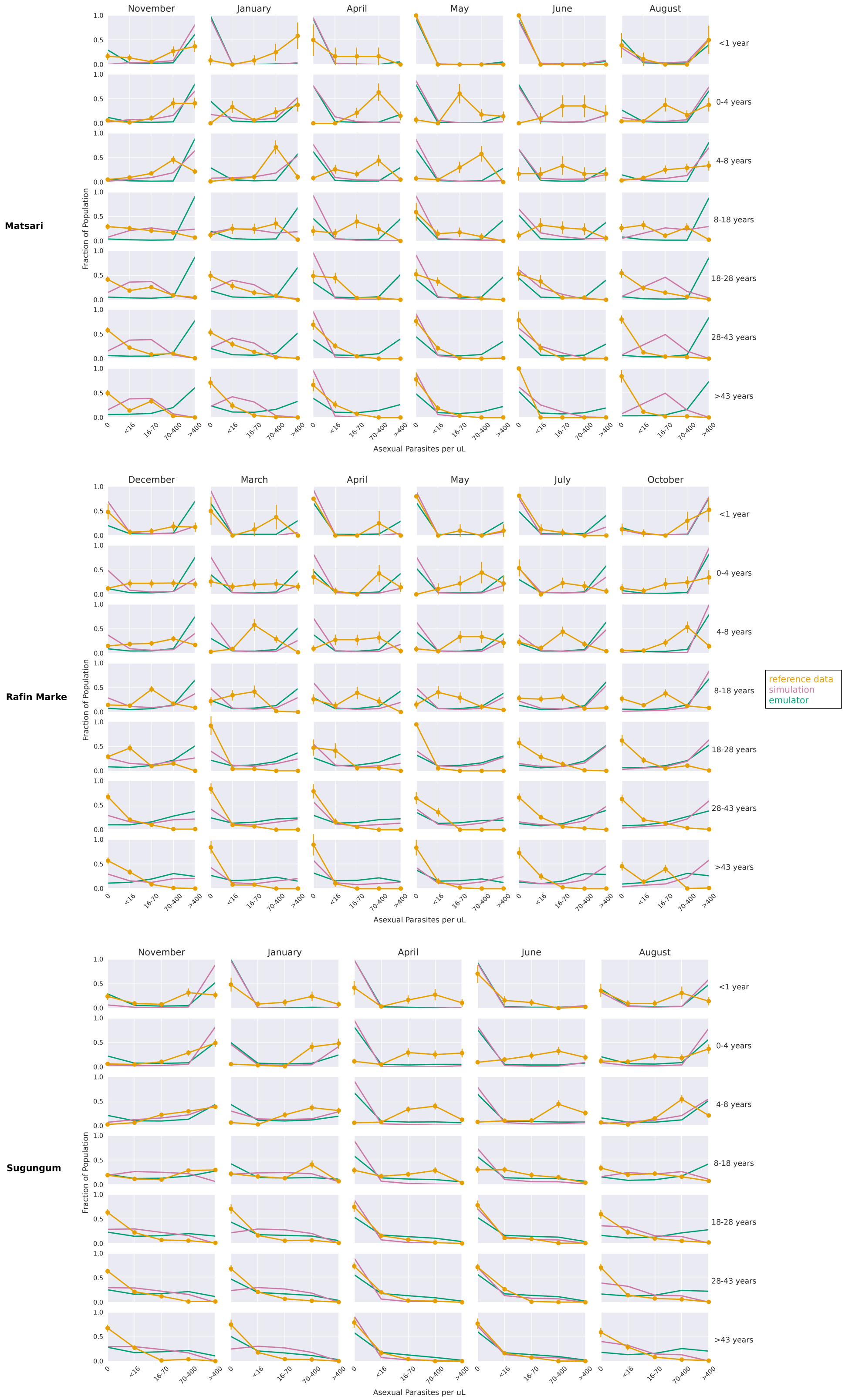


**Fig 3. Site-excluded SGD calibration plots for Matsari, Rafin Marke and Sugungum sites.** These plots show the simulation, emulator, and reference data for the study sites corresponding to the parasitemia outcomes, calibrated via the stochastic gradient descent method, using emulators not explicitly trained on data from these sites.
